# Diverse Strategies Utilized by Coronaviruses to Evade Antiviral Responses and Suppress Pyroptosis

**DOI:** 10.1101/2022.07.29.502014

**Authors:** Xinyu Fu, Weilv Xu, Yang Yang, Danyue Li, Wen Shi, Xinyue Li, Nan Chen, Qian Lv, Yuhua Shi, Jidong Xu, Yuqi Yan, Fushan Shi, Xiaoliang Li

## Abstract

Viral infection triggers inflammasome-mediated caspase-1 activation. Nevertheless, limited understanding exists regarding how viruses use the active caspase-1 to evade host immune response. Here, we use porcine epidemic diarrhea virus (PEDV) as a model of coronaviruses (CoVs) to illustrate the intricate regulation of CoVs to combat IFN-I signaling and pyroptosis. Our findings demonstrate that PEDV infection stabilizes caspase-1 expression via papain-like protease PLP2’s deubiquitinase activity and the enhanced stabilization of caspase-1 disrupts IFN-I signaling by cleaving RIG-I at D189 residue. 6-Thioguanine (6TG), a PLP2 inhibitor, plays a critical role in reversing the inhibitory effect of IFN-I via PLP2 and also inhibiting PEDV replication. Meanwhile, PLP2 can degrade GSDMD-p30 by removing its K27-linked ubiquitin chain at K275 to restrain pyroptosis. Papain-like proteases from other genera of CoVs (PDCoV and SARS-CoV-2) have the similar activity to degrade GSDMD-p30. We further demonstrate that SARS-CoV-2 N protein induced NLRP3 inflammasome activation also uses the active caspase-1 to counter IFN-I signaling by cleaving RIG-I. Therefore, our work unravels a novel antagonistic mechanism employed by CoVs to evade host antiviral response.

## Introduction

Coronaviruses which belong to order Nidovirales, family Coronaviridae, and subfamily Orthocoronavirinae, are a group of viruses packaged in envelop covered with spikes and contain a linear, single and positive-stranded RNA [1]. Coronaviruses are subdivided into four genera, including α, β, γ, and δ. As a member of the α-genus, PEDV infection is characterized by symptoms including vomiting, severe diarrhea, dehydration, and high mortality in suckling piglets [2]. The viral genome of PEDV spans 28 kb and encodes a total of seven open reading frames (ORFs) [3]. Similar to other Coronaviruses (CoVs), PEDV employs its own proteases, specifically the papain-like protease (PLpro) and the main protease, to cleave the polyprotein. The papain-like protease of PEDV is encoded by the largest nonstructural protein Nsp3, which comprises two papain-like protease domains, namely PLP1 and PLP2 or PLpro [4]. In general, α coronaviruses and clade A of β coronaviruses harbor PLP1 and PLP2, and other coronaviruses have one functional PLpro only [5]. Studies have shown that CoVs PLP2 (PLpro) act as a viral deubiquitinase to negatively regulate type I IFN signaling pathway [4, 6]. For example, mouse hepatitis virus (MHV) PLP2 can bind to and deubiquitinate IRF3 to prevent its nuclear translocation [7]. Transmissible gastroenteritis virus (TGEV) PLP2 is able to inhibit the degradation of IκBα by decreasing its ubiquitination, resulting in the suppression of NF-κB signaling [8]. SARS-CoV-2 PLpro can reduce the ubiquitination of RLR signaling, which depends on its deubiquitinating activity [9]. Shin’s study shows that PLPro from SARS-CoV-2 has deISGylase activity for IRF3 and impairs type I interferon response [10]. As to PEDV PLP2, it has been reported that PLP2 deubiquitinates RIG-I and STING to block the IFN signaling pathway [4]. The catalytically dead mutants of PLP2 (C1729A, H1888A and D1901A) can abrogate their deubiquitinase activity and fail to inhibit PEDV-induced IFN-β production.

Caspase-1 is a cysteine protein hydrolase, which participates in cell death and inflammatory response [11]. Caspase-1 exists in the form of proprotein (pro-caspase-1) at static state and acts as an enzyme after activation, which relies on the activation of different inflammasomes [12]. Interestingly, high expression of pro-caspase-1 can also result in its auto-activation in the absence of a ligand [13–15]. Pyroptosis is an inflammatory caspases-dependent, pro-inflammatory programmed cell death characterized by losing of cell membrane integrity, pore forming and swelling and rupture of the cells [16]. The canonical pathway of pyroptosis is mediated by caspase-1 [17, 18]. When host cells’ pattern-recognition receptors (PRRs) are stimulated by pathogen-associated molecular patterns (PAMPs) or damage-associated molecular patterns (DAMPs), inflammasomes will assemble automatically and activate themselves, leading to the activation of caspase-1. Researches have demonstrated that NLRP3 inflammasome can be activated by ZIKV [19], SARS-CoV-2 [20] and MERS-CoV [21, 22], and AIM2 inflammasome can be activated by SARS-CoV-2 [23] while NLRP1 inflammasome can be activated by human rhinovirus (HRV) and picornaviruses [24]. Activated caspase-1 cleaves GSDMD to produce N-terminal GSDMD-p30 fragments with perforating activity on cell membrane, which leads to pyroptosis, and it also cleaves pro-IL-1β to produce IL-1β to improve antiviral capacity [17, 18].

What is clear is that inflammasomes-mediated caspase-1 activation plays a vital role in antiviral defense [25]. Caspase-1-induced pyroptosis, an inflammatory cell death process, has also been proven to have a negative effect on viral replication [26, 27]. Virus-host interactions drive mutual evolution, leading to diversification of viral immune evasion and host antiviral responses. Here, we report that PEDV infection increases the stabilization of pro-caspase-1 by removing its K11-linked ubiquitin chains at K134 via its papain-like protease PLP2. Consequently, caspase-1 targets to RIG-I for cleavage at D189 residue which leads to decreased IFN-I signaling and enhanced PEDV replication. Meanwhile, to assure the integrity of host cells which might be broken by GSDMD-p30 cleaved by caspase-1, PLP2 can degrade GSDMD-p30 by removing its K27-linked ubiquitin chain at K275 residue to restrain pyroptosis. We further demonstrate that papain-like proteases from other genera of CoVs, such as PDCoV and SARS-CoV-2, have the similar activity to degrade GSDMD-p30. Importantly, we confirm that SARS-CoV-2 N protein induced NLRP3 inflammasome activation also uses the active caspase-1 to counter IFN-I signaling. Therefore, our findings uncover a distinctive feature of papain-like protease in antagonizing antiviral responses that might serve as a target for CoVs treatment in the future.

## Results

### PEDV infection elevates the expression of caspase-1

In order to investigate the effect of PEDV on porcine caspase-1, endogenous caspase-1 expression was analyzed by immunoblotting in intestinal tissues of non-infected and PEDV naturally-infected piglets. The results showed that expression of caspase-1 was increased in the PEDV-infected piglets (Fig 1A). qRT-PCR showed that *caspase-1* mRNA abundance was not altered in IPEC-J2 cells upon PEDV infection (Fig 1B). Immunoblotting also confirmed that PEDV infection promoted the amount of caspase-1 in IPEC-J2 cells and CHX treatment further elevated caspase-1 expression (Fig 1C and S1A Fig), suggesting that the increased caspase-1 expression was not caused by the transcription of mRNA and synthesis of new proteins. In addition, Vero cells were transfected with plasmids encoding caspase-1 and then infected with PEDV. Caspase-1 was found to be upregulated during PEDV infection with varying MOI and infectious time (Fig 1D and 1E). We further demonstrated that overexpression of caspase-1 alone resulted in active caspase-1 p20 (Fig 1F), which is consistent with previous reports that the high expression of caspsae-1 causes auto-activation and produces active caspase-1 p20 [13–15]. Since active caspase-1 p20 can lead to the cleavage of GSDMD, we next investigated the relationship between PEDV infection and pyroptosis. Vero cells were co-transfected with GSDMD-FL (full length) and caspase-1 and then infected with PEDV. Caspase-1 expression and cleavage of GSDMD increased progressively in both MOI and time dependent manner during infection (Fig 1G and 1H). Interestingly, PEDV infection resulted in declines in LDH release and propidium iodide (PI) uptaking compared to non-infected group, despite the increased GSDMD cleavage (Fig 1I, 1J and S2 Fig). These results indicate that PEDV infection promotes the amount of caspase-1.

**Fig 1.**
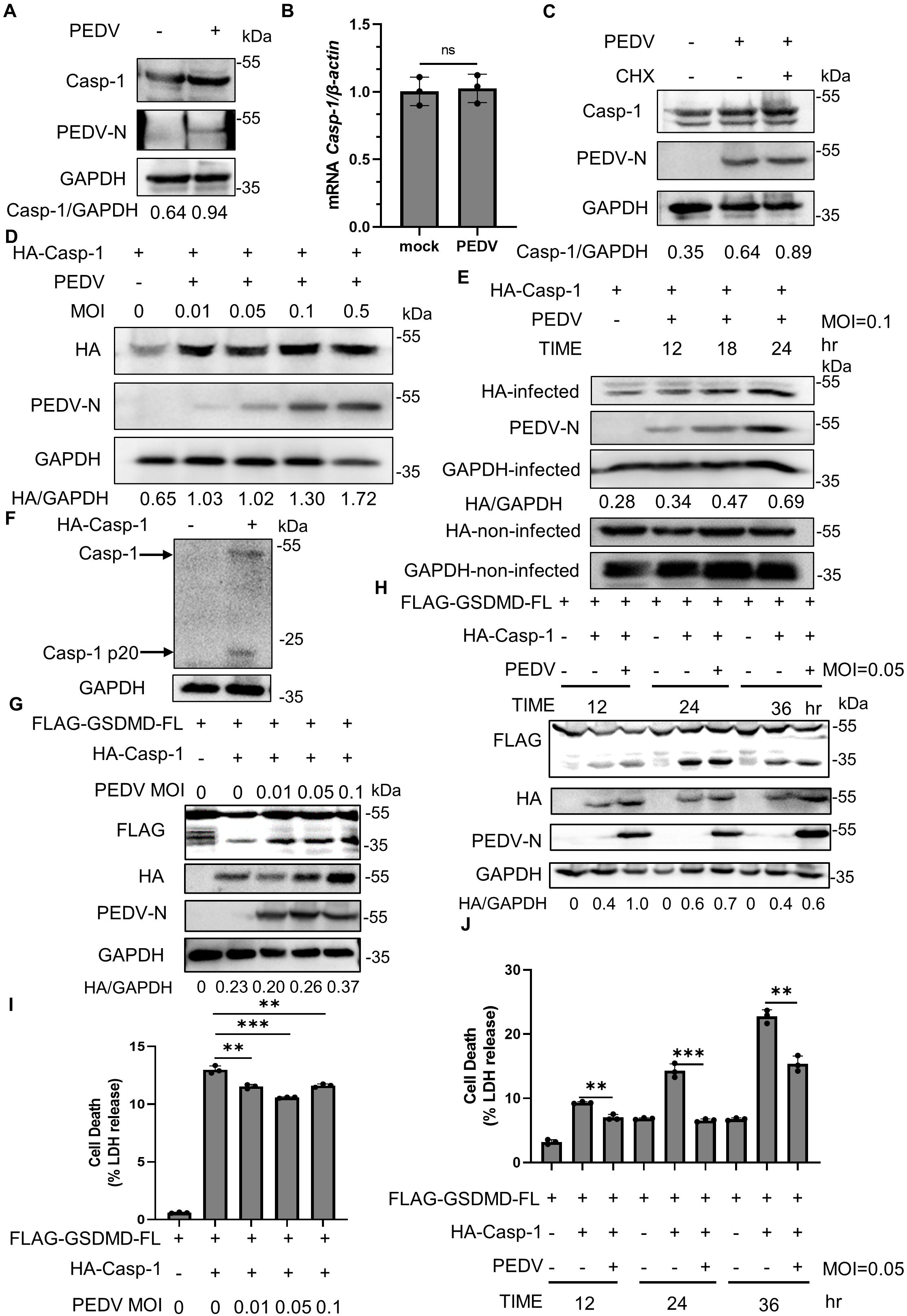
PEDV infection elevates the expression of caspase-1. (A) The intestinal tissue lysates were extracted from healthy or PEDV naturally-infected piglets for immunoblotting (IB). (B and C) IPEC-J2 cells were non-infected or infected with PEDV at an MOI of 0.5, followed by cycloheximide (CHX) treatment (25 μg/ml) for 6 h. The indicated gene mRNA was quantified by qRT-PCR (B). The indicated proteins were analyzed by immunoblotting (C). (D) Vero cells were transfected with plasmid encoding porcine caspase-1. At 24 h after transfection, the cells were non-infected or infected with different doses of PEDV for another 24 h, the cell lysates were then processed for immunoblotting. (E) Vero cells were transfected with plasmid encoding porcine caspase-1. At 24 h after transfection, the cells were non-infected or infected with PEDV at an MOI of 0.1. At indicated time points afterinfection, the cell lysates were analyzed by immunoblotting. (F) HEK293T cells were transfected with plasmid encoding caspase-1. At 24 h after transfection, the cell lysates were analyzed by immunoblotting. (G and I) Vero cells were co-transfected with plasmids encoding GSDMD-FL and caspase-1. At 12 h after transfection, the cells were non-infected or infected with different doses of PEDV for another 24 h, the cell lysates were processed for immunoblotting (G), the supernatants were collected for LDH release assay (I). (H and J) Vero cells were co-transfected with plasmids encoding GSDMD-FL and caspase-1. At 12 h after transfection, the cells were non-infected or infected with PEDV at an MOI of 0.05. At indicated time points after infection, the cell lysates were analyzed by immunoblotting (H), the supernatants were collected for LDH release assay (J). All results shown are representative of at least three independent experiments. ****stands for P<0.0001, ***stands for P<0.001, ** stands for P<0.01, * stands for P<0.05 and ns stands for non-significant difference.

### The papain-like protease 2 of PEDV inhibits the proteasomal degradation of caspase-1

The increased expression of caspase-1 induces GSDMD cleavage but inhibits pyroptosis. Therefore, we speculated that some kind of viral proteins affected this process. Vero cells transfected with GSDMD-FL were infected with PEDV, following performed with coimmunoprecipitation (Co-IP) and mass spectrometry (MS) analysis. The papain-like protease 2 (PLP2) of PEDV by marker peptide “KVELDATK” was found (S1B Fig). The Co-IP assay also confirmed that PEDV PLP2 interacted with GSDMD-FL (S1C Fig). Since it is known that GSDMD-FL and caspase-1 are interacted [28], we further examined whether PLP2 interacts with caspase-1. HEK293T cells were co-transfected with FLAG-tagged caspase-1 and MYC-tagged PLP2, following treated with VX765, an inhibitor of caspase-1, to curb caspase-1 activation. Co-IP assay showed that PLP2 interacted with caspase-1 (S1D Fig). Indirect immunofluorescence further demonstrated that GSDMD, caspase-1 and PEDV PLP2 were colocalized in the cytoplasm (S1E Fig). Thus, PLP2 interacts with both caspase-1 and GSDMD.

Given that high expression of caspase-1 leads to auto-activation, caspase-1-C285A (caspase-1 mutant, which lacks its protease activity) and PLP2 were co-transfected into HEK293T cells. As shown in Fig 2A, we observed an increased protein amount of caspase-1-C285A in the presence of PLP2. In addition, co-transfection of PLP2, GSDMD-FL and caspase-1 in HEK293T also revealed that PLP2 caused the accumulation of caspase-1 but not GSDMD-FL (S3A Fig). We next measured caspase-1 degradation by a cycloheximide (CHX) chase assay to determine whether PLP2 could stabilize caspase-1. The results demonstrated that PEDV PLP2 could stabilize caspase-1 but not GSDMD-FL by delaying its degradation (Fig 2B and S3B Fig). Ubiquitination is a key signal for proteasomal degradation, and PLP2 has the deubiquitinase activity. To identify whether the deubiquitinase activity of PLP2 is required for the stabilization of caspase-1, we generated the catalytic mutants (C113A, H272A, D285A) of PLP2 and co-transfected them with caspase-1-C285A. As expected, all of the three mutants lost the ability to stabilize caspase-1 (Fig 2C). Furthermore, we co-expressed the wild type PLP2 or its mutants with HA-tagged ubiquitin (HA-Ub) and FLAG-tagged caspase-1 in HEK293T cells. Compared to the empty vector, wild type PLP2 substantially inhibited ubiquitination of caspase-1 while the three mutants had no effect on it (Fig 2D), suggesting that the deubiquitinase activity of PLP2 is indispensable for the stabilization of caspase-1.

**Fig 2.**
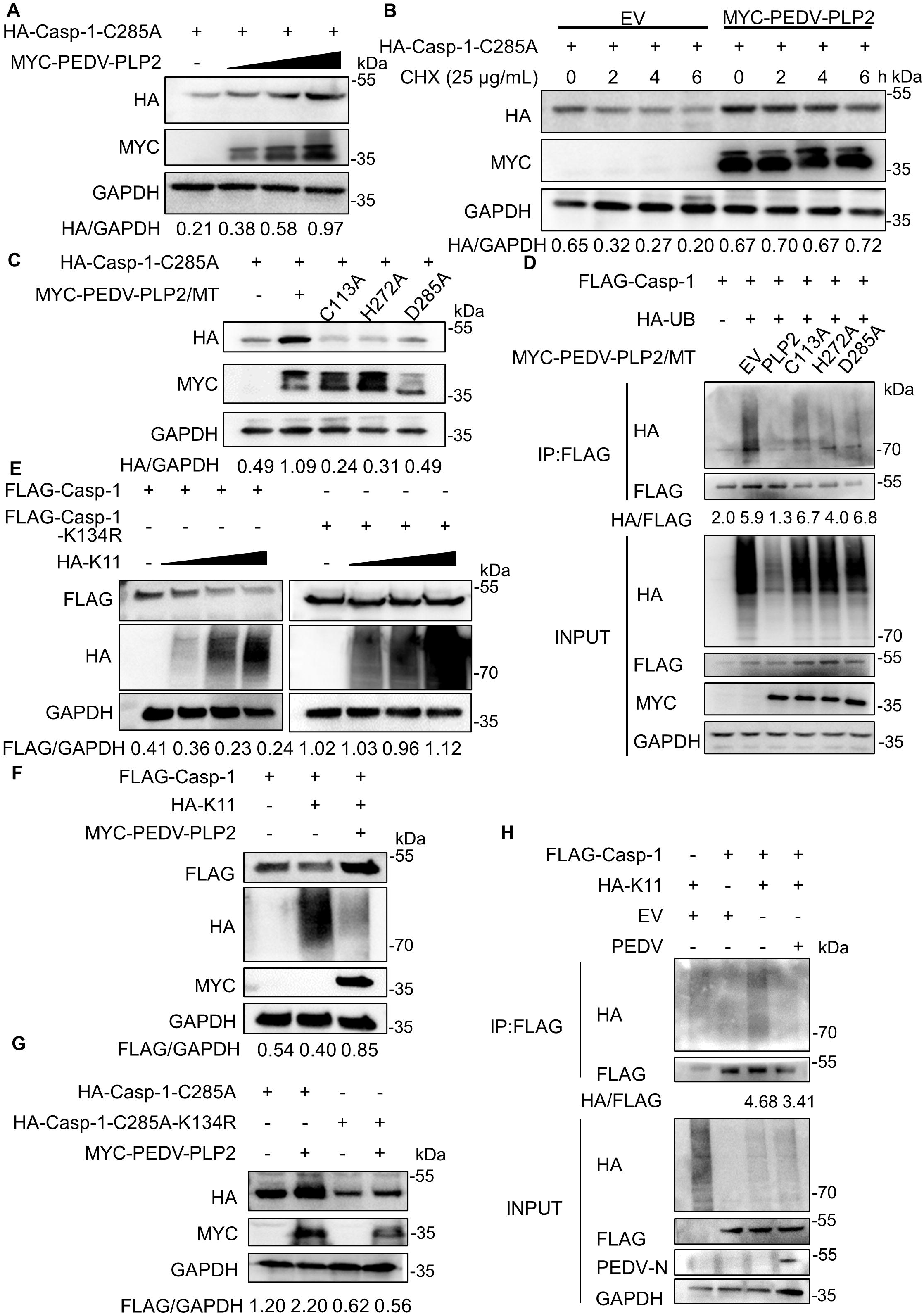
PLP2 inhibits the proteasomal degradation of caspase-1 by removing K11-linked ubiquitin chains at Lys134. (A) HEK293T cells were co-transfected with HA-caspase-1-C285A and increasing amount of MYC-PEDV-PLP2. Cell lysates were analyzed by immunoblotting. (B) HEK293T cells were co-transfected with HA-caspase-1-C285A together with empty vector or MYC-PEDV-PLP2 then treated with cycloheximide (CHX) (25 μg/ml) for the indicated time points. Cell lysates were analyzed by immunoblotting. (C) HEK293T cells were co-transfected with HA-caspase-1-C285A together with wild-type PEDV PLP2 or its protease-defective mutants (C113A, H272A and D285A). Cell lysates were analyzed by immunoblotting. (D) Immunoprecipitation analysis of HEK293T cells expressing FLAG-caspase-1 and HA-UB together with MYC-tagged empty vector, wild-type PEDV-PLP2 or its mutants as indicated. Anti-FLAG immunoprecipitates were analyzed by immunoblotting (IP). The expression levels of the transfected proteins were analyzed by immunoblotting (INPUT). (E) HEK293T cells were co-transfected with FLAG-caspase-1 or its mutant FLAG-caspase-1-K134R together with increasing amount of HA-tagged K11-linked ubiquitin (HA-K11). Cell lysates were analyzed by immunoblotting. (F) HEK293T cells were transfected with FLAG-caspase-1 together with HA-K11 and MYC-PEDV-PLP2 as indicated. Cell lysates were analyzed by immunoblotting. (G) HEK293T cells were transfected with HA-caspase-1-C285A or its mutant HA-caspase-1-C285A-K134R together with MYC-tagged empty vector or PEDV PLP2. Cell lysates were analyzed by immunoblotting. (H) Vero cells were transfected with FLAG-caspase-1 together with HA-K11 or empty vector. At 24 h after transfection, the cells were non-infected or infected with PEDV at an MOI of 0.05 for another 24 h. Immunoprecipitation was performed with anti-FLAG binding beads and analyzed by immunoblotting. All results shown are representative of at least three independent experiments.

It has been reported that the K11-linked ubiquitination of human caspase-1 at Lys-134 is a way to induce its degradation [29]. To determine whether K134 residue of porcine-derived caspase-1 is also critical for its ubiquitination, we constructed porcine-derived caspase-1-K134R mutant in which the lysine (K) residue was replaced with arginine (R). As shown in Fig 2E, wild type caspase-1 was degraded by K11 in a dose-dependent manner, while caspase-1-K134R could not be degraded by K11. Moreover, we found that caspase-1-K134R mutant exhibited a dramatic decrease in K11-linked ubiquitination (S3C Fig). To further confirm the role of PLP2 in this process, we co-expressed PLP2 with caspase-1 and K11 ubiquitin, the results demonstrated that the degradation effect of K11 on caspase-1 was abrogated by PLP2 (Fig 2F). Immunoblotting also showed that PLP2 stabilized caspase-1 but not caspase-1-K134R (Fig 2G). Additionally, PEDV infection decreased K11-linked poly-ubiquitination of caspase-1, further fortifying the above findings (Fig 2H). Taken together, these results indicate that PLP2 of PEDV stabilizes porcine caspase-1 by removing K11-linked ubiquitin chains at Lys134.

### Caspase-1 attenuates IFN-I signaling during PEDV infection

It has been reported that caspase-1 can negatively regulate IFN-I signaling by cleaving cGAS during DNA virus infection [30]. As an RNA virus, PEDV activates IFN-I signaling through the RLRs pathway. Since IFN-I plays a vital role in antiviral process, we wondered whether the stabilized caspase-1 can affect it. We initially found that overexpression of caspase-1 in IPEC-J2 cells attenuated the *IFN-α*, *IFN-β*, *ISG15* and *OAS1* mRNA levels induced by poly (I:C) treatment (Fig 3A-3D). In addition, caspase-1 inhibitor VX765 reversed the attenuated IFN-I signaling during poly (I:C) treatment (Fig 3E and 3F). Moreover, knockout of caspase-1 in THP-1 cells further promoted the expression of *IFN-β* and *ISG15* mRNA induced by poly (I:C) (Fig 3G and 3H). Same results were observed for PLP2 in IPEC-J2 cells (Fig 3I and 3J). Thus, we determined that both caspase-1 and PLP2 were antagonists of IFN-I. Next, the above-mentioned findings were further verified in PEDV-infected IPEC-J2 cells, which were consistent with our observations (Fig 3K-3P). These results indicate that PEDV-induced IFN-I signaling was counteracted by the stabilized caspase-1.

**Fig 3.**
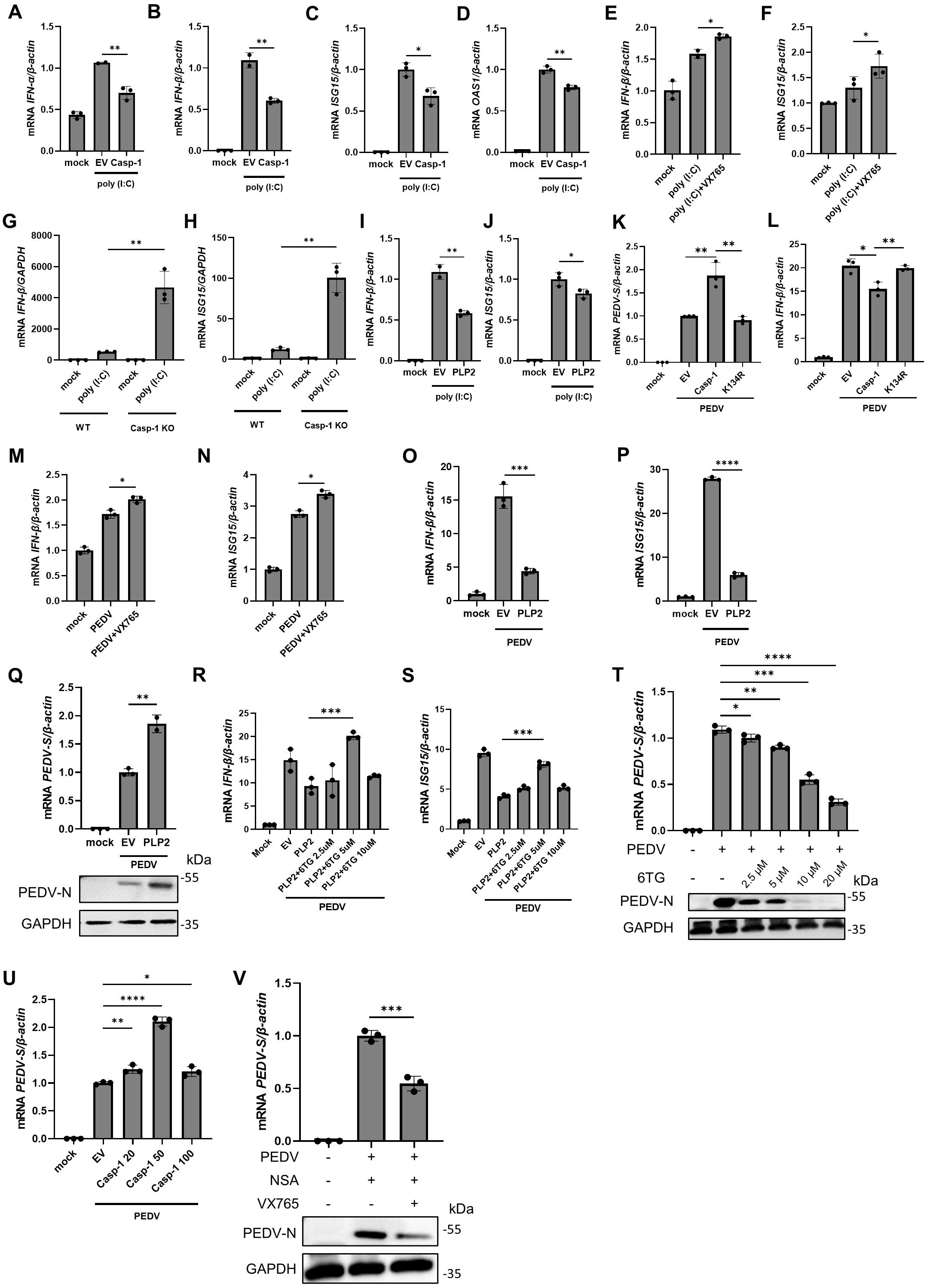
Caspase-1 attenuates IFN-I signaling during PEDV infection. (A-D) Relative qRT-PCR analysis of *IFN-α*, *IFN-β*, *ISG15* and *OAS1* mRNA levels in IPEC-J2 cells transfected with empty vector or caspase-1 for 24 h, and then stimulated with poly (I:C) (200 ng/ml) for another 12 h. (E and F) Relative qRT-PCR analysis of *IFN-β* and *ISG15* mRNA levels in IPEC-J2 cells treated with VX765 (20 μM) or DMSO, followed by poly (I:C) (200 ng/ml) transfection for 12 h. (G and H) Relative qRT-PCR analysis of *IFN-β* and *ISG15* mRNA levels in wild type (WT) or caspase-1 knockout (KO) THP-1 cells transfected with poly (I:C) (200 ng/ml) for 12 h. (I and J) Relative qRT-PCR analysis of *IFN-β* and *ISG15* mRNA levels in IPEC-J2 cells transfected with empty vector or PEDV PLP2 for 24 h, followed with poly (I:C) (200 ng/ml) transfection for another 12 h. (K and L) Relative qRT-PCR analysis of *PEDV S* and *IFN-β* mRNA levels in IPEC-J2 cells transfected with empty vector or caspase-1 or caspase-1-K134R for 12 h, followed with PEDV infection (MOI=0.5) for another 24 h. (M and N) Relative qRT-PCR analysis of *IFN-β* and *ISG15* mRNA levels in IPEC-J2 cells treated with VX765 (20 μM) or DMSO, followed with PEDV infection (MOI=0.5) for 24 h. (O and P) Relative qRT-PCR analysis of *IFN-β* and *ISG15* mRNA levels in IPEC-J2 cells transfected with empty vector or PEDV PLP2 for 24 h, followed with PEDV infection (MOI=0.5) for another 24 h. (Q) IPEC-J2 cells were transfected with empty vector or PEDV PLP2 for 24 h, followed with PEDV infection at an MOI of 0.5 for another 24 h. The indicated gene mRNA levels were quantified by qRT-PCR and the indicated proteins were analyzed by immunoblotting. (R and S) Relative qRT-PCR analysis of *IFN-β* and *ISG15* mRNA levels in IPEC-J2 cells transfected with empty vector or PEDV PLP2 for 24 h, followed with PEDV infection at an MOI of 0.5 and different concentrations of 6TG for anitoher 24 h. (T) IPEC-J2 cells were treated with different concentrations of 6TG followed by PEDV infection at an MOI of 0.5 for 24 h. The indicated gene mRNA levels were quantified by qRT-PCR and the indicated proteins were analyzed by immunoblotting. (U) Relative qRT-PCR analysis of *PEDV S* mRNA levels in IPEC-J2 cells transfected with empty vector or increasing amount of caspase-1 for 12 h, followed with PEDV infection (MOI=0.1) for another 24 h. (V) IPEC-J2 cells were treated with NSA and/or VX765 followed by PEDV infection at an MOI of 0.5 for 24 h. The indicated gene mRNA levels were quantified by qRT-PCR and the indicated proteins were analyzed by immunoblotting. All results shown are representative of at least three independent experiments. ****stands for P<0.0001, ***stands for P<0.001, ** stands for P<0.01, stands for P<0.05 and ns stands for non-significant difference.

The role of caspase-1 and PLP2 in viral replication was further analyzed considering the vital antiviral effect of IFN-I. PLP2 was transfected into IPEC-J2 cells and then infected with PEDV. Consistent with its antagonistic effect on IFN-I, the results showed that PLP2 could promote PEDV replication, which was also confirmed by immunoblotting (Fig 3Q). The small molecule compound 6-Thioguanine (6TG) has been identified as an inhibitor of various coronavirus papain-like proteases. Therefore, we used 6TG for further validation and found that it can reverse the inhibition of IFN-I by PLP2 and also inhibit PEDV replication (Fig 3R-3T). Since the high expression of caspase-1 leads to autoprocessing, we transfected caspase-1 with different doses and then treated with PEDV infection. As shown in Fig 3U, all doses of caspase-1 could promote PEDV replication, while the effect of high concentration of caspase-1 was attenuated, which might relate to pyroptosis caused by caspase-1 activation. Conversely, the inhibitor VX765 effectively inhibits PEDV replication, which was also confirmed through immunoblotting (Fig 3V). Together, these data suggest that the enhanced caspase-1 stabilized by PEDV infection attenuates type I IFN signaling and thus benefits viral replication. Specific inhibitors, however, effectively target viral and host proteins, thus inhibiting viral replication.

### Cleavage of RIG-I by caspase-1 ensures PEDV replication

As caspase-1 is vital to suppress type I IFN during PEDV infection, we hypothesized that caspase-1 exerted its effect on RIG-I. Therefore, we analyzed whether caspase-1 could directly interact with RIG-I. HEK293T cells were then transfected with MYC-tagged RIG-I and FLAG-tagged caspase-1, Co-IP assay showed that RIG-I interacted with caspase-1 (Fig 4A). Interestingly, we found that RIG-I was diminished after co-transfected with caspase-1 (Fig 4B). Thus, we co-transfected caspase-1 or its mutant with RIG-I at different doses in HEK293T, followed with immunoblotting. As shown in Fig 4C, a clear band with a molecular weight around 25-35 kDa was observed, while the caspase-1-C285A (caspase-1 mutant) impaired the cleavage. Because RIG-I was N-terminal MYC-tagged, we speculated that caspase-1 cut RIG-I at N-terminus. We therefore generated mutants of RIG-I at different sites to identify the exact cleavage site of RIG-I by caspase-1. RIG-I or its mutants (D163A, D189A, D194A, D199/202/203A and D209A) were co-transfected with caspase-1 in HEK293T cells. As shown in Fig 4D and S4A Fig, wild-type RIG-I, D163A, D194A, D199/202/203A and D209A could still be cleaved by caspase-1 successfully, while RIG-I-D189A could not be cleaved. These results imply that RIG-I was a cleaved target of caspase-1 at D189. Moreover, we investigated the relationship between human derived caspase-1 and RIG-I. As shown in S4B Fig, a band with a molecular weight around 25-35 kDa was observed (lane 2). Based on the multiple-sequence alignment of RIG-Is (S4C Fig), different mutants, including single or double mutants of human RIG-I (h-RIG-I) were generated and co-expressed with empty vector or h-caspase-1. The results showed that the cleaved fragments from all of the mutants were reduced compared to the wild type RIG-I (S4B Fig). We next generated a triple mutant of RIG-I (h-RIG-I-D163/194/234A) and transfected it with caspase-1 in HEK293T cells. S4D Fig revealed that the cleaved band between 25-35 kDa was further weakened, suggesting that the D163, D194 and D234 residues of RIG-I might be the cleavage sites of human derived caspase-1.

**Fig 4.**
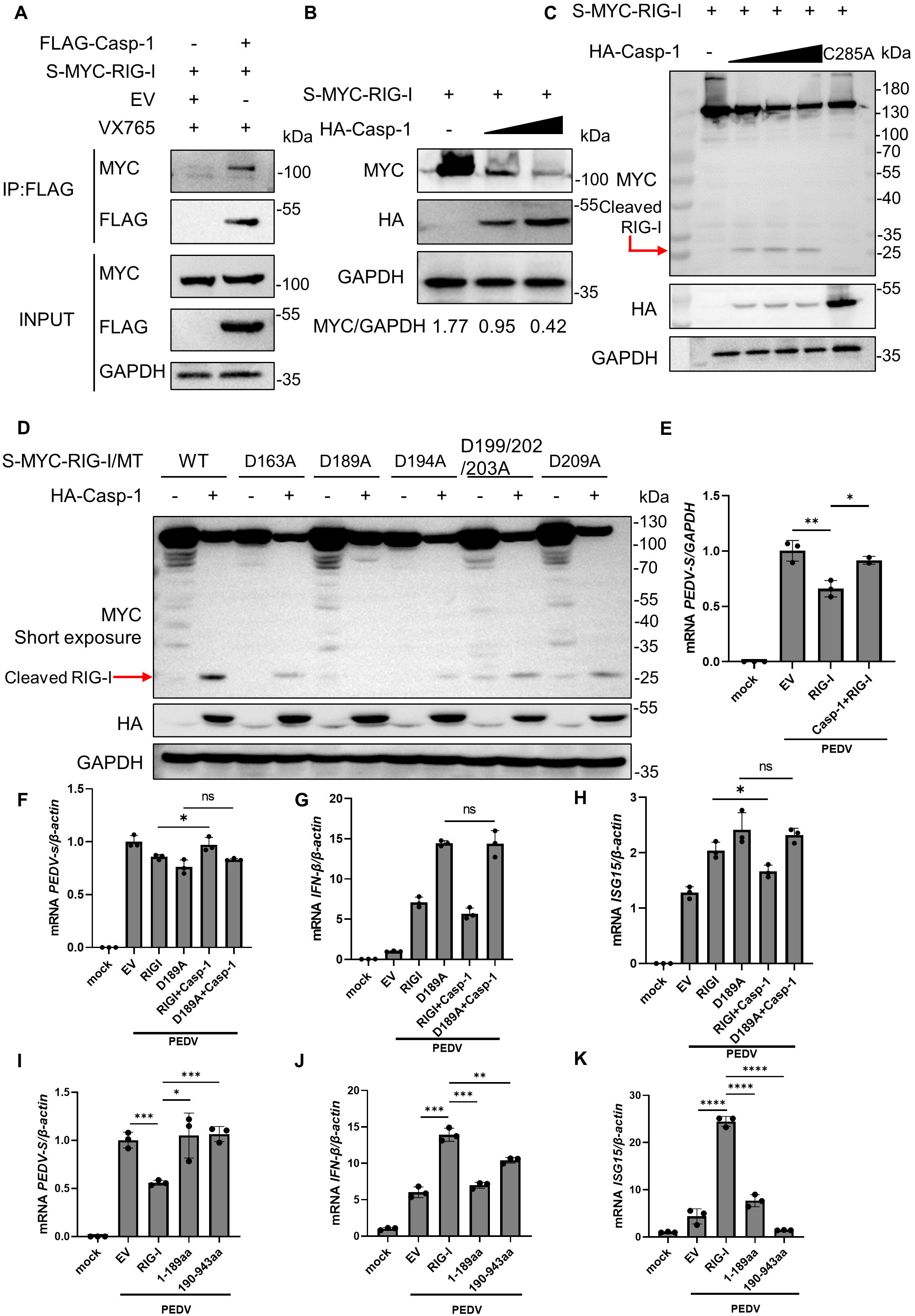
Cleavage of RIG-I by caspase-1 ensures PEDV replication. (A) HEK293T cells co-transfected with FLAG-caspase-1 and S-MYC-RIG-I (porcine derived) were lysed and immunoprecipitated with anti-FLAG beads and analyzed by immunoblotting. (B and C) HEK293T cells were co-transfected with S-MYC-RIG-I and increasing amount of HA-caspase-1 or HA-caspase-1-C285A. Cell lysates were analyzed by immunoblotting. (D) HEK293T cells were co-transfected with wild-type (WT) S-MYC-RIG-I or its mutants (D163A, D189A, D194A, D199/202/203A, D209A) together with empty vector or HA-caspase-1. Cell lysates were analyzed by immunoblotting. (E) Vero cells were transfected with empty vector, RIG-I together with or without caspase-1. At 12 h after transfection, the cells were non-infected or infected with PEDV at an MOI of 0.05. The indicated gene mRNA levels were quantified by qRT-PCR. (F-H) Relative qRT-PCR analysis of *PEDV S, IFN-β* and *ISG15* mRNA levels in IPEC-J2 cells transfected with empty vector, RIG-I, RIG-I-D189A, RIG-I and Caspase-1 or RIG-I-D189A and Caspase1 for 24 h, followed by PEDV infection (MOI=0.5) for another 24 h. (I-K) Relative qRT-PCR analysis of *PEDV S, IFN-β* and *ISG15* mRNA levels in IPEC-J2 cells transfected with empty vector, RIG-I, RIG-I-1-189aa or RIG-I-190-943aa for 24 h, followed by PEDV infection (MOI=0.5) for another 24 h. All results shown are representative of at least three independent experiments. ****stands for P<0.0001, ***stands for P<0.001, ** stands for P<0.01, * stands for P<0.05 and ns stands for non-significant difference.

To further investigate the effects of RIG-I cleavage on viral replication and type I IFN signaling, Vero cells were transfected with empty vector (EV), RIG-I or co-transfected with caspase-1 and RIG-I, followed by PEDV infection. As shown in Fig 4E, RIG-I significantly attenuated viral replication, while the co-transfected group reversed this effect. We then generated a RIG-I mutant, RIG-I-D189A, and transfected into IPEC-J2 cells and then infected with PEDV. As expected, wild type RIG-I facilitated *IFN-β* and *ISG15* production and inhibited PEDV replication, and the RIG-I-D189A mutant further enhanced IFN-I signaling and impaired viral replication. Furthermore, the simultaneous expression of RIG-I and caspase-1 resulted in a decrease in IFN-β levels and an increase in PEDV replication. Conversely, co-transfection of RIG-I-D189A and caspase-1 yielded outcomes comparable to the expression of RIG-I-D189A (Fig 4F-4H). We transfected IPEC-J2 cells with either RIG-I-WT or RIG-I-D189A, followed by a 12-hour poly(I:C) stimulation. The results showed that the RIG-I-D189A mutant exhibited increased production of IFN-β and ISG15 transcripts in response to poly(I:C), resembling the response observed during PEDV infection (S4E and S4F Fig). This phenomenon is likely due to the presence of endogenous caspase-1 in the cells, which has the capacity to cleave RIG-I. However, because of the limited amount of endogenous caspase-1 in the absence of PLP2’s stabilizing function, the cleavage is not as pronounced as observed during PEDV infection.

Next, we evaluated the function of RIG-I-1-189aa and RIG-I-190-943aa in activating the innate immune response as well as in viral replication. We found that wild-type RIG-I promoted IFN induction and inhibited PEDV replication, while RIG-I-1-189aa or RIG-I-190-943aa potently reversed IFN-I production and restored viral reproduction (Fig 4I-4K). Taken together, these results collectively demonstrate that caspase-1 targets the D189 residue of porcine RIG-I for cleavage and the cleaved fragments lose their function to inhibit PEDV replication.

### PLP2 degrades GSDMD-p30 by removing the K27-linked ubiquitin chains at K275 residue to inhibit pyroptosis

The activation of caspase-1 can lead to GSDMD cleavage and thus cause pyroptotic cell death. However, according to the above results, PEDV infection did not cause substantial pyroptosis at early stage (Fig 1H and 1I). To determine whether PLP2 also targets GSDMD and regulates pyroptosis, HEK293T cells were initially co-transfected with GSDMD-FL and caspase-1 for 6 h, and then PLP2 was transfected in a dose-dependent manner. The immunoblotting results showed that the cleaved fragment of GSDMD (GSDMD-p30) was decreased in the presence of PLP2 (Fig 5A). We also conducted additional experiments, including groups of caspase-1-C285A, which is unable to cleave GSDMD, and PLP2-C113A, which lacks enzymatic activity and cannot inhibit pyroptosis or degrade GSDMD-p30 (S5 Fig). Meanwhile, we repeatedly observed a reduced protein level of GSDMD-p30 after overexpression of PLP2, but had no variation on GSDMD-FL (Fig 5B and S6A Fig). The LDH release assay and the propidium iodide staining also confirmed that PLP2 had an inhibitory effect on pyroptotic cell death induced by GSDMD-p30 (Fig 5C and S6B Fig). As shown in Fig. 5D, PLP2 had a stronger interaction with GSDMD-p30 than GSDMD-FL, which further demonstrated the complex relationship between GSDMD-p30 and PLP2. In order to evaluate the effect of PLP2 on GSDMD-p30 more directly, we co-transfected GFP-tagged PLP2 and FLAG-tagged GSDMD-p30 in HEK293T cells. GSDMD-p30 was found to localize and formed bright red clumps in the cytoplasm and causing cell membrane disruption (Fig 5E, top row). In the presence of PLP2, GSDMD-p30 was evenly dispersed in cytoplasm, no swelling or discontinuity in the membrane is observed (Fig 5E, bottom row). Taken together, we reveal that PLP2 degrades GSDMD-p30 and thus blocks pyroptosis.

**Fig 5.**
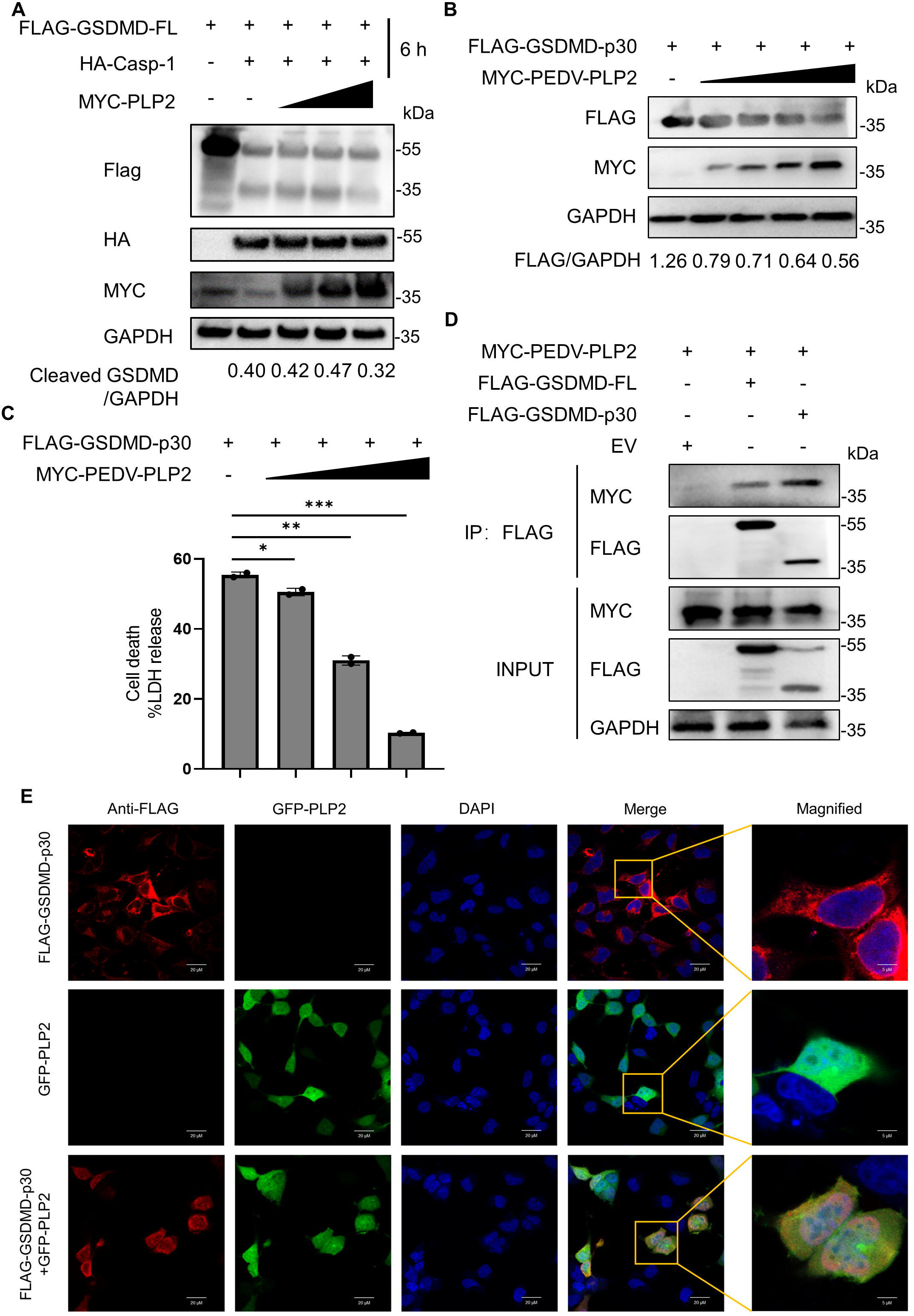
PLP2 degrades GSDMD-p30 to inhibit pyroptosis. (A) HEK293T cells were initially co-transfected with FLAG-GSDMD-FL and HA-caspase-1. At 6 h after transfection, the cells were transfected with increasing amount of MYC-PEDV-PLP2. Cell lysates were analyzed by immunoblotting. (B and C) HEK293T cells were co-transfected with FLAG-GSDMD-p30 and increasing amount of MYC-PEDV-PLP2. Cell lysates were analyzed by immunoblotting (B). The supernatants were collected for LDH release assay (C). (D) HEK293T cells co-transfected with MYC-PEDV-PLP2 together with empty vector, FLAG-GSDMD-FL or FLAG-GSDMD-p30 were lysed and immunoprecipitated with anti-FLAG beads and analyzed by immunoblotting. (E) HEK293T cells were transfected with FLAG-GSDMD-p30 and GFP-PLP2 respectively or together for 24 h, and then FLAG-GSDMD-p30 cells were labeled with indicated antibodies. Subcellular localization of FLAG-GSDMD-p30 (red), GFP-PLP2 (green), and DAPI (blue, nucleus marker) were visualized with confocal microscopy. All results shown are representative of at least three independent experiments. ****stands for P<0.0001, ***stands for P<0.001, ** stands for P<0.01, * stands for P<0.05 and ns stands for non-significant difference.

To determine which degradation system is responsible for the degradation of GSDMD-p30, we examined the effect of the proteasome inhibitor MG132, the autophagy inhibitor 3MA and CQ on GSDMD-p30 degradation in the presence of PLP2. The results showed that PLP2-mediated degradation of GSDMD-p30 was rescued by MG132. While it was previously documented that GSDMD-p30 is unstable and prone to degradation via the proteasome when the pore formation does not occur, our results suggest a connection between GSDMD-p30 degradation induced by PLP2 and the proteasome pathway (Fig 6A). We further revealed that all the catalytic mutants of PLP2 (C113A, H272A, D285A) lost the ability to degrade GSDMD-p30 which was also confirmed by the LDH release assay (Fig 6B and 6C), suggesting that the deubiquitinase activity of PLP2 is essential for GSDMD-p30 degradation. To determine the lysine residues of GSDMD-p30 to which ubiquitin is attached, we used the UbPred program (http://www.ubpred.org), to predict the potential ubiquitination sites of porcine GSDMD-p30. This analysis revealed three putative lysine residues: K103, K177 and K275. We then generated GSDMD-p30-K103R, K177R and K275R mutants in which each of these lysine residues was replaced with arginine (R). We co-transfected the wild type porcine GSDMD-p30 and three of its mutants with HA-tagged ubiquitin, followed with immunoblotting. As shown in Fig 6D, co-transfection of HA-Ub had no effect on the expression of GSDMD-p30-K275R mutant, suggesting that Lys275 residue might be the ubiquitination site of GSDMD-p30. Next, we co-transfected HA-tagged ubiquitin mutants (with only one of the seven lysine residues retained as lysine, while the other six replaced with arginine) with GSDMD-p30 in HEK293T cells. The LDH release assay revealed that K27-linked ubiquitin mutant markedly elevated the LDH level (Fig 6E), which was consistent with our previous study [31]. Ubiquitination assay further confirmed that K27-linked ubiquitin promoted the ubiquitination of GSDMD-p30 (Fig 6F, lane 2 and 4) and K275R mutant displayed a significant reduction of K27-linked ubiquitination (Fig 6F, lane 4 and 5). Moreover, we co-expressed the wild type PLP2 or its mutants with HA-tagged K27-linked ubiquitin and FLAG-tagged GSDMD-p30 and then performed Co-IP assay. As expected, wild type PLP2 dramatically reduced K27-linked ubiquitination of GSDMD-p30, while neither of its mutants performed the same effect (Fig 6G). Taken together, our results demonstrate that PLP2 is able to degrade GSDMD-p30 by removing the K27-linked ubiquitin chain at K275 residue of GSDMD-p30 and thus abrogate pyroptosis.

**Fig 6.**
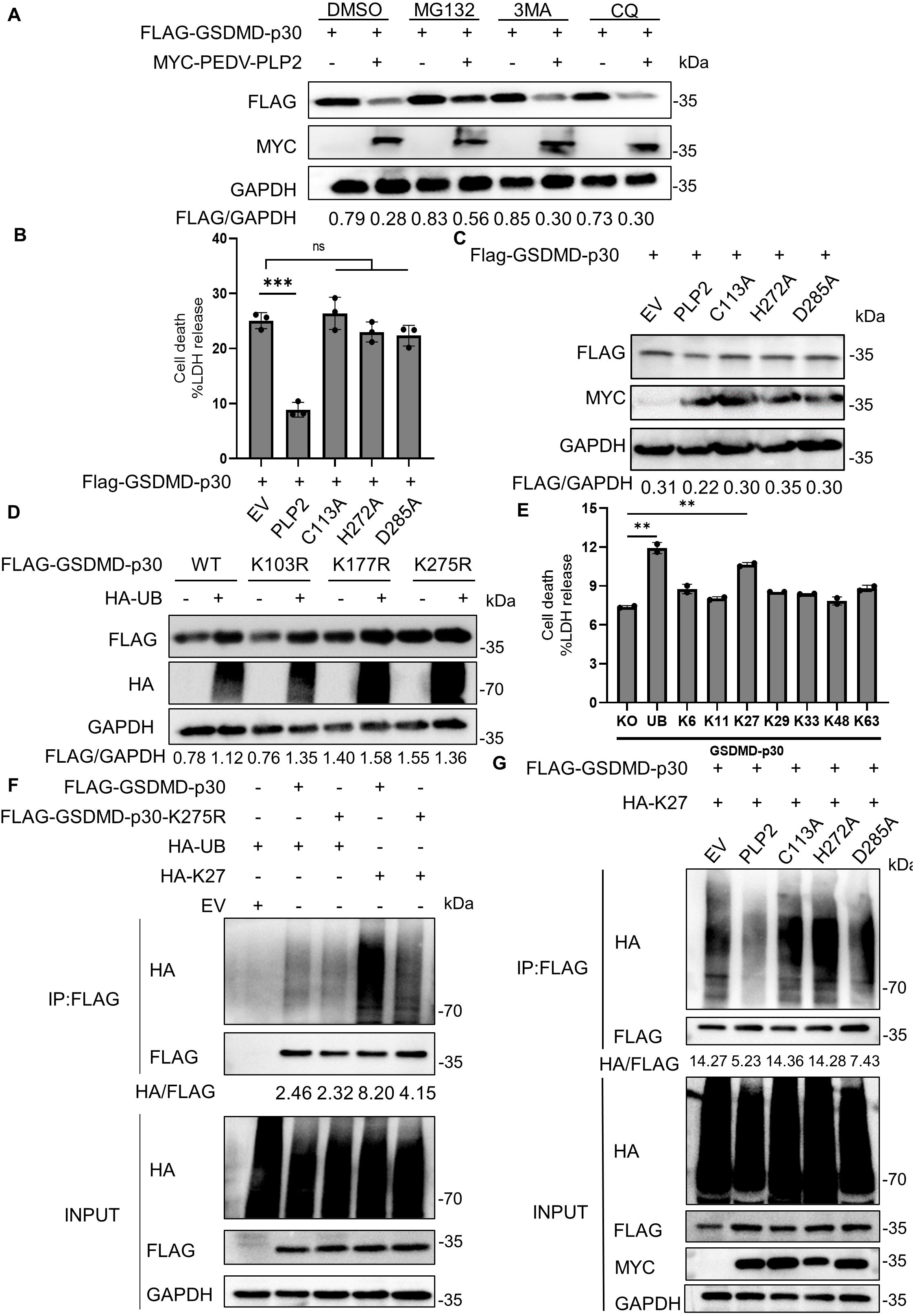
PLP2 removes the K27-linked ubiquitin chain at K275 residue of GSDMD-p30. (A) HEK293T cells were transfected with FLAG-GSDMD-p30 together with empty vector or MYC-PEDV-PLP2 then treated with DMSO, MG132 (10 μM), 3MA (1 mM) or CQ (40 μM) for 6 h. Cell lysates were analyzed by immunoblotting. (B and C) HEK293T cells were co-transfected with FLAG-GSDMD-p30 together with empty vector, wild type PLP2 or its protease-defective mutants (C113A, H272A, D285A). The supernatants were collected for LDH release assay (B). Cell lysates were analyzed by immunoblotting (C). (D) HEK293T cells were co-expressed with FLAG-GSDMD-p30 or its mutants (K103R, K177R, K275R) together with empty vector or HA-UB. Cell lysates were analyzed by immunoblotting. (E) HEK293T cells were co-transfected with FLAG-GSDMD-p30 and HA-tagged WT ubiquitin or its mutants (K6, K11, K27, K29, K33, K48, K63, KO). The supernatants were collected for LDH release assay. (F) Co-immunoprecipitation and immunoblot analysis of extracts of HEK293T cells transfected with FLAG-GSDMD-p30 or FLAG-GSDMD-p30-K275R together with HA-UB or HA-K27 ubiquitin. (G) Co-immunoprecipitation and immunoblot analysis of extracts of HEK293T cells transfected with FLAG-GSDMD-p30 and HA-K27 ubiquitin together with empty vector, wild type PLP2 or its mutants (C113A, H272A, D285A). All results shown are representative of at least three independent experiments. ****stands for P<0.0001, ***stands for P<0.001, ** stands for P<0.01, stands for P<0.05 and ns stands for non-significant difference.

### CoVs utilize different strategies to disrupt antiviral response and inhibit pyroptosis

Next, we tested whether papain-like proteases in other genus of CoVs have the same effects on GSDMD-p30 and caspase-1. PEDV-PLP2, SARS-CoV-2-PLpro and PDCoV-PLpro were generated and then transfected into HEK293T cells with GSDMD-p30. As shown in Fig 7A, 7B and S7A Fig, all these papain-like proteases could inhibit LDH release, PI uptake and degrade GSDMD-p30, which revealed their ability in abrogating pyroptosis. Also, these papain-like proteases had no discernible impact on GSDMD-FL (S7B Fig). Furthermore, we examined the role of these viral proteins in stabilizing caspase-1 and found that only PEDV PLP2 could promote caspase-1 expression (Fig 7C). It is possible that PDCoV and SARS-CoV-2 use different ways to activate caspase-1. To evaluate the effect of these viral proteins on PEDV replication, IPEC-J2 cells were transfected with empty vector or these papain-like proteases, and then infected with PEDV. All of these papain-like proteases exhibited the potential to enhance viral replication (Fig 7D). Collectively, GSDMD-p30 was attenuated by papain-like proteases of CoVs, which led to higher viral replication.

**Fig 7.**
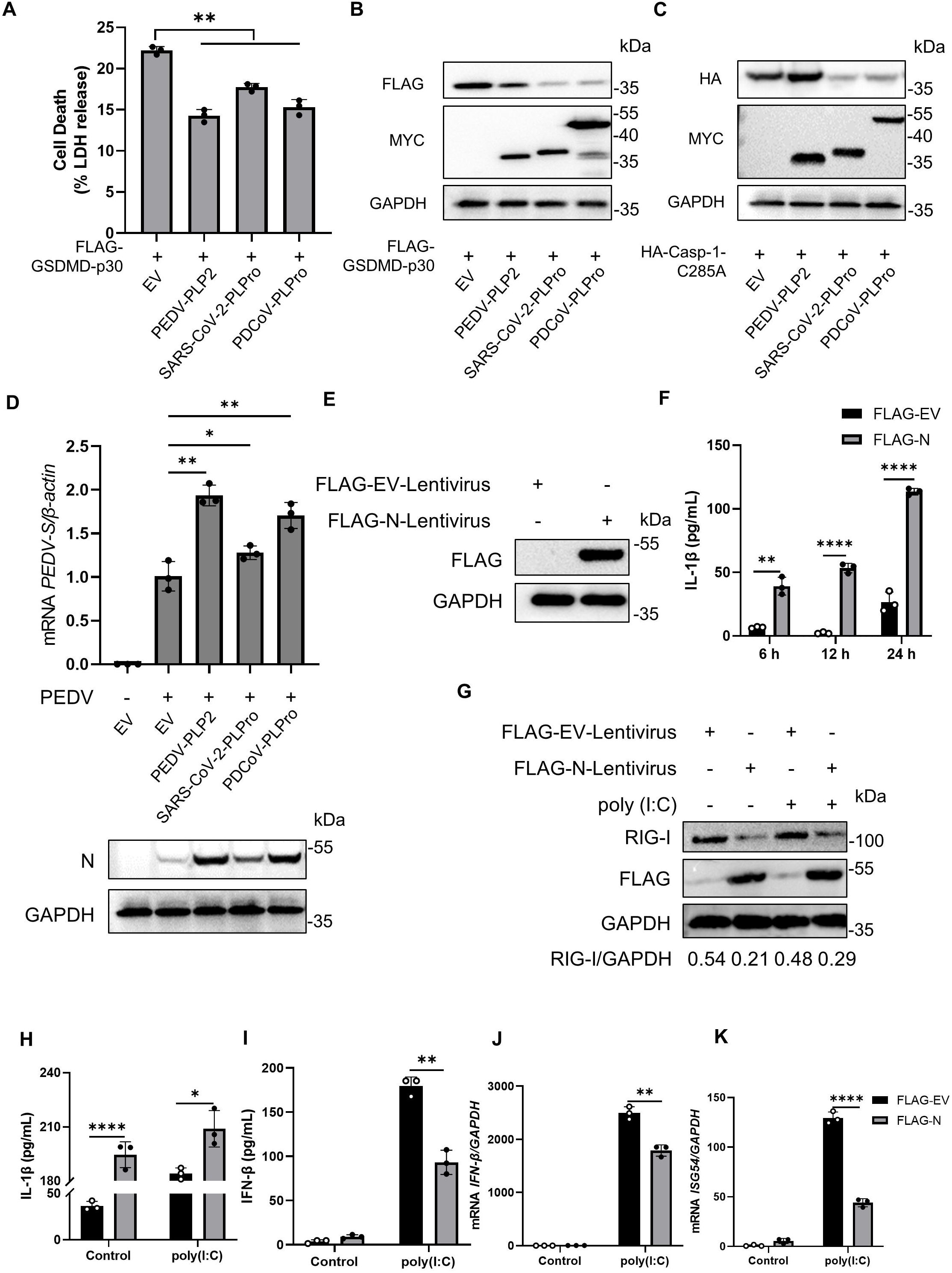
Papain-like proteases in other CoVs can also abrogate pyroptosis by degradation of GSDMD-p30. (A and B) HEK293T cells were transfected with FLAG-GSDMD-p30 together with empty vector, PEDV-PLP2, SARS-CoV-2-PLPro or PDCoV-PLPro. The supernatants were collected for LDH release assay (A). Cell lysates were analyzed by immunoblotting (B). (C) HEK293T cells were transfected with HA-caspase-1-C285A together with empty vector, PEDV-PLP2, SARS-CoV-2-PLPro or PDCoV-PLPro. Cell lysates were analyzed by immunoblotting. (D) IPEC-J2 cells were transfected with empty vector, PEDV-PLP2, SARS-CoV-2-PLPro or PDCoV-PLPro. At 24 h after transfection, the cells were infected with PEDV at an MOI of 0.1 for another 24 h. The indicated gene mRNA levels were quantified by qRT-PCR (top). Cell lysates were analyzed by immunoblotting (bottom). (E) Immunoblot analysis of extracts of THP-1 cells stably infected with FLAG-EV-Lentivirus or FLAG-N-Lentivirus. (F) PMA-differentiated THP-1 macrophages were stably infected with FLAG-EV-Lentivirus or FLAG-N-Lentivirus. The supernatants were collected and analyzed by ELISA for IL-1β. (G-L) PMA-differentiated THP-1 macrophages were stably infected with FLAG-EV-Lentivirus or FLAG-N-Lentivirus, followed with poly (I:C) (200 ng/ml) transfection for 6 h. Cell lysates were analyzed by immunoblotting (G). The supernatants were collected and analyzed by ELISA for IL-1β (H) and IFN-β (I) assay. The mRNA levels of *IFN-β* (J) and *ISG54* (K) were quantified by qRT-PCR. All results shown are representative of at least three independent experiments. ****stands for P<0.0001, ***stands for P<0.001, ** stands for P<0.01, * stands for P<0.05 and ns stands for non-significant difference.

It has been reported that the nucleocapsid protein (N) of SARS-CoV-2 activates the NLRP3 inflammasome, thus we hypothesized that NLRP3 inflammasome-mediated caspase-1 activation could also antagonize IFN production. In that case, we generated THP-1 cell lines stably expressing FLAG-EV-Lentivirus (negative control) or FLAG-N-Lentivirus (Lentivirus carrying the SARS-CoV-2 N gene) (Fig 7E). Consistent with previous studies [20], FLAG-N-Lentivirus elevated the secretion of IL-1β, suggesting that SARS-CoV-2 N can activate NLRP3 inflammasome (Fig 7F). To evaluate the role of SARS-CoV-2 N in IFN production, the differentiated macrophages were then stimulated with poly (I:C) and we found that the endogenous RIG-I was significantly reduced in the presence of SARS-CoV-2 N (Fig 7G). As shown in Fig 7H and 7I, the production of IL-1β was elevated while the IFN-β was dramatically reduced by SARS-CoV-2 N. Moreover, qRT-PCR analyses revealed that the expression of *IFN-β* and *ISG54* mRNA level were notably repressed by SARS-CoV-2 N (Fig 7J and 7K), indicating that the activated caspase-1 targeted RIG-I to abrogate IFN-I signaling. Taken together, we unravel a novel antagonistic mechanism employed by different viral proteins from CoVs and thus evade host IFN signaling and pyroptosis (Fig 8).

**Fig 8.**
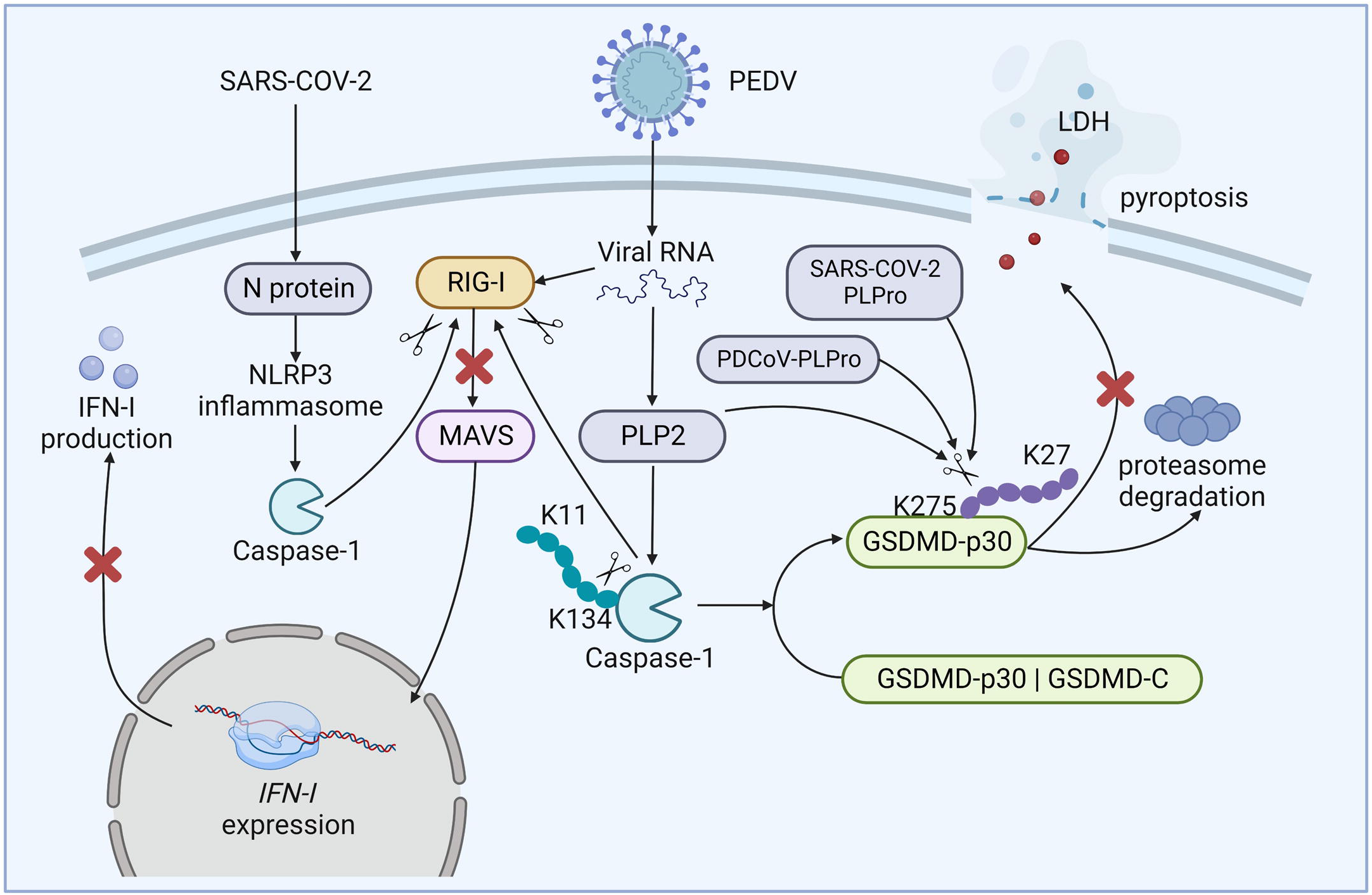
Mechanistic diagram of CoVs antagonizing antiviral responses and pyroptosis.

## Discussion

A variety of viruses have been reported to activate inflammasomes. The inflammasome serves as a platform for caspase-1 activation, leading to caspase-1-dependent IL-1β release and subsequent pyroptosis. This, in turn, increases the expression of antiviral genes to control invading pathogens [32]. As a countermeasure, viruses have evolved strategies to antagonize host immune response. Previous study has demonstrated that DNA virus-induced inflammasome-mediated active caspase-1 cleaves human cGAS at D140 and D157 to dampen IFN production [30]. In the present study, we elucidated the intricate regulation of RNA virus to counteract IFN-I signaling and pyroptosis using active caspase-1 and papain-like protease. We found that CoVs-induced active caspase-1 disrupts IFN-I signaling by cleaving RIG-I to produce inactive fragments. Meanwhile, papain-like protease of CoVs can degrade GSDMD-p30 by removing its K27-linked ubiquitin chain at K275 residue to restrain pyroptosis. Therefore, our work unravels a novel antagonistic mechanism employed by CoVs to evade host antiviral response.

To date, most viruses induced caspase-1 activation is mediated by inflammasomes. EV71 3D and ZIKV NS5 activate the NLRP3 inflammasome by interacting with NACHT and the LRR domain of NLRP3 [19, 33, 34]. NS1 of ZIKV also enhances NLRP3 inflammasome activation to facilitate its infection [29]. The influenza A virus (IAV) non-structural protein PB1-F2 contributes to severe pathophysiology through triggering NLRP3-dependent caspase-1 activation [35]. Except for NLRP3, viruses can also induce other inflammasomes activation. Human rhinovirus (HRV) 3C protease directly cleaves human NLRP1 between the Glu130 and Gly131 junction, which liberates the activating C-terminal fragment and subsequently promoting NLRP1 inflammasome activation [36]. NLRP9b recognizes short double-stranded RNA stretches via RNA helicase Dhx9 and forms inflammasome complexes with ASC and pro-caspase-1 to promote caspase-1-dependent IL-1β release and pyroptosis [37]. Herpes simplex virus 1 (HSV1) can induce AIM2-, pyrin- and ZBP1-mediated caspase-1 activation, cytokine release and cell death [38]. For coronaviruses, studies have shown that MERS-CoV and at least three different proteins of SARS-CoV-2 can induce NLRP3 inflammasome activation and caspase-1-mediated subsequent pyroptosis [20, 22, 39, 40]. Evidence also demonstrated that circulating monocytes from COVID-19 patients show signs of AIM2 inflammasome activation and caspase-1-meidated GSDMD cleavage and pyroptosis [23]. Unlike the reported manner of caspase-1 activation, our results suggest that PEDV induces porcine caspase-1 activation via its papain-like protease PLP2 through elevating pro-caspase-1 expression by removing its K11-linked ubiquitin chains at K134 residue. Studies have demonstrated that accumulated pro-caspase-1 lead to autoprocessing within the catalytic domain [13–15]. Consequently, active caspase-1 targets to porcine RIG-I for cleavage at D189 residue leading to decreased IFN-I signaling and enhanced PEDV replication. Interestingly, PLPro from SARS-CoV-2 and PDCoV did not use this way to activate caspase-1, as they can trigger NLRP3- or AIM2-mediated caspase-1 activation. Similarly, inflammasome-mediated caspase-1 activation also cleaved human RIG-I to abrogate IFN-I signaling pathway. Therefore, our study suggests that the main purpose of virus-induced inflammasome-mediated caspase-1 activation is to antagonize IFN-I signaling. A newly published study further indicated that SARS-CoV-2 infection induced high expression of pro-caspase-4/11, which may result in its auto-activation in the absence of a ligand [41]. Although our study is limited at the protein level, it uncovers a new and different mechanism of CoVs inducing caspase-1 activation, which provides a new idea for further study in this area.

Activation of inflammasome is often vital in the host antiviral immune response [40, 42–44]. It is obvious that triggering of pyroptosis and inflammasomes activation is detrimental to the replication and survival of viruses. Thus, viruses have also developed strategies to antagonize extensive inflammasome activation and pyroptosis, especially at the early stage of viral infection. The human papillomavirus (HPV) oncoprotein E7 promotes the TRIM21-mediated K33-linked ubiquitination of the IFI16 inflammasome for degradation to inhibit pyroptosis. African swine fever virus (ASFV) pS273R cleaves GSDMD at G107-A108 to generate a shorter N-terminal of GSDMD (GSDMD-N_1-107_) which is not capable of triggering pyroptosis. The accessory protein (PB1-F2) of H5N1 and H3N2 influenza A viruses (IAV) can bind to the pyrin and LRR domains of NLRP3 and prevent NEK7 binding to stabilize the auto-repressed and closed conformation of NLRP3. 2A and 3C proteases of EV71 can cleave NLRP3 directly to inhibit the activation of inflammasome, while the EV71 3C-like protease can also cleave GSDMD to block the pyroptosis pathway. Our recent research also proved that 3C-like proteases Nsp5 of coronaviruses (PEDV, PDCoV, SARS-CoV-2 and MERS-CoV) can cleave GSDMD at Q193 to produce two fragments that are unable to trigger pyroptosis. SARS-CoV-2 N induces pro-inflammatory cytokines through promoting the assembly and activation of the NLRP3 inflammasome [20]. However, SARS-CoV-2 N protein is also capable of binding to GSDMD directly to inhibit the GSDMD cleavage from caspase-1 to avoid pyroptosis [45]. Our results demonstrated that SARS-CoV-2 N-induced NLRP3 inflammasome activation can antagonize IFN production through cleaving RIG-I by activated caspase-1. Furthermore, as depicted in Figure 4D, we noticed that the cleavage bands for the D163A, D194A, D199/202/203A, and D209A mutants exhibited reduced intensity when compared to the WT RIG-I, implying that D189 may not be the sole cleavage site. Moreover, it is plausible that this phenomenon is subject to regulation by additional factors. Consequently, further comprehensive research is warranted to unveil the underlying mechanisms. In the present study, we further demonstrated that CoVs (PEDV, PDCoV and SARS-CoV-2) can abrogate pyroptosis by degradation of GSDMD-p30 via their papain-like proteases. Therefore, we discovered a novel mechanism for CoVs to antagonize pyroptosis. Nevertheless, our findings were constrained by technological and instrumental limitations since they were contingent upon the overexpression of plasmids in cells. This necessitates further investigation.

In conclusion, we used PEDV as a model of CoVs to illustrate the sophisticated regulation of CoVs to counteract IFN-I signaling and pyroptosis. For the first time, we demonstrate the role of inflammasome dependent or independent caspase-1 activation in antagonizing IFN-I signaling. Furthermore, we show that papain-like protease of CoVs can inhibit pyroptosis by degradation of GSDMD-p30 through proteasome pathway. Thus, our present study unveils a novel antagonistic mechanism by which CoVs manipulates the cross-talk of IFN signaling and pyroptosis, which might indicate a framework for design of anti-CoVs therapies.

## Materials and methods

### Reagents and antibodies

Anti-HA (3724) and anti-Caspase-1 (2225) antibodies were purchased from Cell Signaling Technology. Anti-FLAG antibody (F1804), anti-MYC antibody (C3956) and anti-FLAG magnetic beads (10004D) were obtained from Sigma. Anti-FLAG antibody (rabbit source), Goat pAb to MS IgG (Chromeo 546, ab60316), Dnk pAb to Rb IgG (Alexa Fluor 647, ab150075) fluorescent secondary antibodies were acquired from Abcam. Anti-GAPDH antibody, HRP-labeled goat anti-rabbit IgG and goat anti-mouse IgG were from Hangzhou Fudebio. Anti-GSDMDC1 antibody (sc-393581) was purchased from Santa Cruz. Anti-RIG-I antibody (20566) was obtained from proteintech. The anti-PEDV N monoclonal antibody was prepared in our laboratory as previously described [46]. Efficient eukaryotic transfection reagent VigoFect was obtained from Vigorous Biotechnology (Beijing), Lipo8000 transfection reagent was obtained from Beyotime Biotechnology while Lipofectamine 2000 transfection reagent was obtained from Invitrogen.

### Plasmids

Eukaryotic expression vectors used in this subject were saved in our laboratory. The PLP2 fragment sequence was amplified from cDNA of PEDV strain ZJ15XS0101 (GenBank accession no. KX55SO0281) which was isolated by our collaborating laboratories for generational preservation, while amplification of the porcine GSDMD gene and the porcine caspase-1 gene used cDNA of IPEC-J2 as a templet. The PLP2 gene was cloned to PRK-MYC vector, and the porcine GSDMD gene was cloned onto p3×FLAG-CMV vector. The porcine caspase-1 gene was cloned to PCMV-HA and p3×FLAG-CMV vector, respectively. Porcine GSDMD-p30 fragment was amplified from the p3×FLAG-N-GSDMD-FL plasmid and cloned onto p3×FLAG-CMV. PLP2, caspase-1 and GSDMD-p30 enzyme active site mutants were constructed based on the eukaryotic expression plasmids PRK-MYC-PEDV-PLP2, HA-PCMV-Caspase-1 and p3×FLAG-GSDMD-p30. The plasmids encoding SARS-CoV-2 PLPro, PDCoV PLPro and SARS-CoV-2-N were synthesized by Tsingke Biotech. Primers used in the plasmids construction are listed in S1 Table.

### Cell culture and transfection

IPEC-J2 cells, Vero cells and HEK293T cells were cultured in Dulbecco’s modified Eagle’s medium (DMEM) with added 10% FBS and 1% penicillin-streptomycin. Wild type THP-1 and caspase-1 deficient THP-1 were cultured in RPMI 1640 medium containing 10% FBS and 1% penicillin-streptomycin solution. The PEDV strain ZJ15XS0101 (GenBank accession no. KX55SO0281) was isolated and stored in our laboratory [47]. When HEK293T cells seeded in plates grow to about a density of 60%∼80%, indicated plasmids were transfected to the cells by VigoFect according to the manufacturer’s operation guide. When IPEC-J2 and Vero seeded in plates grow to approximately 70%∼80%, they were transfected with corresponding plasmids using Lipo8000 transfection reagent according to the manufacturer’s introduction. Lipo2000 was applied in the transfection of poly(I:C) (Merck) to IPEC-J2 during the second transfection period. All animal studies were reviewed and approved by the Animal Care and Use Committee of Zhejiang University.

### Viral infection

Viral infection was performed when the IPEC-J2 or Vero cells grow to the density around 80%∼90% or after the transfection of other indicated plasmids.

### Lentivirus infection

*SARS-COV-2-N* over-expressing THP-1 cells were generated using lentiviral infection technique. After cloning *SARS-COV-2-N* coding sequence to the pLVX-IRES-Puro-3×Flag lentiviral vector plasmid, HEK293T cells were co-transfected with pLVX-IRES-Puro-3×Flag-SARS-COV-2-N, pMD2G and PSPAX2. Supernatant was collected after 48 h incubation for further lentiviral concentration.

### Inhibitors treatment

The caspase-1 specific inhibitor VX765 diluted to 20 μM was used after plasmid transfection, during viral infection or before poly(I:C) transfection. Protein synthesis inhibitor cycloheximide (CHX) diluted to 25 μg/ml was added at different time points before receiving samples. 6TG, diluted to different concentrations was used at the time of the viral infection. Pyroptosis inhibitor NSA was diluted to 8 μM and performed on indicated cells. Fresh culture medium containing proteasome inhibitor MG132, autophagy initiation inhibitor 3MA or autophagy lysosome inhibitor CQ of appropriate concentration were added to different wells of cell-culturing plates 6 h before receiving samples.

### LDH assay

The culture medium from cells transfected with the corresponding plasmids for 24 hours or infected for the indicated durations was collected. LDH release was assessed using the Cytotoxicity Detection Kit (Promega) following the manufacturer’s instructions. Optical density (OD) values were measured at 492 nm using a microplate reader (Thermo Scientific).

### Immunoblotting

Total proteins were extracted from cells lysed by RIPA lysis buffer (Beyotime Biotechnology) supplemented with 1% Phenylmethanesulfonyl fluoride (PMSF) (Beyotime Biotechnology). After being separated on the 10% SDS-PAGE gel (Verde Biotechnology), protein stripes were transferred onto the polyvinylidene difluoride (PVDF) membranes (Bio-rad). Membranes were blocked in the blocking buffer (Beyotime Biotechnology), followed by incubation with primary antibodies and positioning with secondary antibodies. Chemiluminescent signals were captured by an ECL chemiluminescence imaging analysis system (Clinx Science Instruments).

### LC-MS/MS analysis

Vero cells transfected with GSDMD-FL were infected with PEDV, following performed with coimmunoprecipitation. Immunoprecipitants prepared from whole-cell lysates or gel-filtrated fractions were resolved on SDS-PAGE gels, and protein bands were excised. The samples were digested with trypsin, and then subject to LC-MS/MS analysis.

### Co-immunoprecipitation

Cells were lysed using IP lysis buffer (Beyotime Biotechnology) for 15 min at 4[, and then centrifuged at speed 10000 rpm at 4 [for 15 min. After being incubated with the supernatant of the cell lysis solution in a flip mixer (Kylin-Bell) overnight, the magnetic beads coated with FLAG-antibodies (Sigma) were washed with coIP-buffer for 5 times. Soaking in 1×loading buffer, magnetic beads absorbed with target proteins were denatured for 10 min in a boiling water bath. After discarding magnetic beads, the remaining protein samples were applied to subsequent immunoblotting assay.

### Confocal immunofluorescence assay

HEK293T cells were seeded on glass slides placed in 24-well plates. After being transfected for 24 h, cells were fixed with Immunol Staining Fix Solution (Beyotime), permeabilized with Immunostaining Permeabilization Solution with Saponin (Beyotime), blocked with QuickBlock Blocking Buffer for Immunol Staining (Beyotime), and then incubated with indicated antibodies. The cells were observed under a laser confocal microscope (Olympus).

### ELISA

Supernatants collected from transfected cells were applied to the detection of IL-1β and IFN-β according to the manufacturer’s instructions. Each trial group was conducted independently for three times.

### Total RNA extraction and reverse transcription

Add RNA-easy Isolation Reagent (Vazyme Biotechnology) to cells to gather lysis solution containing cells. Follow the manufacturer’s introduction to extract the RNA. After measuring the product RNA concentration, use the HiScript III RT SuperMix for qPCR (+gDNA wiper) (Vazyme Biotechnology) to reverse transcribe RNA to cDNA.

### Quantitative real-time polymerase chain reaction (qRT-PCR)

qRT-PCR was performed with ChamQ Universal SYBR qPCR Master Mix (Vazyme Biotechnology) according to the manufacturer’s requirements. Primers demanded in this analysis were listed in S2 Table.

### RNA interference

SiRNAs (Genepharma) specific for porcine GSDMD were transfected into IPEC-J2 cells using the Lipofectamine 8000 transfection reagent according to the manufacturer’s instructions. The sequences of GSDMD siRNAs were listed in S3 Table.

### Propidium iodide staining

Cells were incubated with Propidium iodide (PI) (BD Bioscience) for 15 min under light-proof conditions after transfection with indicated plasmids for 24 h. Dyeing condition was observed under fluorescence microscope (Nikon).

### Statistical analysis

All experiments were performed independently at least three times. Data were presented as the mean ± standard deviation (SD), analyzed and used for statistical graphing by GraphPad Prism 8, the significance of differences was determined by two-way ANOVA or Student’s *t*-test. The significance of differences ranked as: ****stands for P<0.0001, ***stands for P<0.001, ** stands for P<0.01, * stands for P<0.05 and ns stands for non-significant difference.

## Acknowledgements

We thank Dr. Ying Shan in the Shared Experimental Platform for Core Instruments, College of Animal Sciences, Zhejiang University for assistance with analysis of laser confocal microscopy imaging.

## Author Contributions

X.L. and F.S. conceived the study. X.F., W.X. and Y.Y. performed most of the experiments. D.L., X.L., N.C., Q.L., Y.S., W.S. and J.X. performed some experiments. X.F., W.X., X.L. and Y.Yan contributed to data analysis. X.F., W.X., Y.Y., J.X. and F.S. contributed to writing the manuscript.

## Data Availability

The data that support the findings of this study are available from the corresponding author upon reasonable request.

## Supplementary Materials

Figs. S1 to S7

Tables S1 to S3

## Supplementary Materials

**S1 Fig. PEDV PLP2 interacts with both caspase-1 and GSDMD.** (A) Grayscale analysis of WB bands in Fig 1C. (B) Vero cells were transfected with FLAG-GSDMD-FL and then infected with PEDV at an MOI of 0.05. Immunoprecipitation was performed with anti-FLAG antibody. The potential GSDMD-binding viral proteins were evaluated using MS analysis. The papain-like protease 2 (PLP2) of PEDV was identified by the marker peptide “KVELDATK”. (C) Co-immunoprecipitation and immunoblot analysis of extracts of HEK293T cells transfected with MYC-PEDV-PLP2 together with FLAG-GSDMD-FL or FLAG-tagged empty vector. (D) Co-immunoprecipitation and immunoblot analysis of extracts of HEK293T cells transfected with MYC-PEDV-PLP2 together with FLAG-caspase-1 or FLAG-tagged empty vector, then treated with VX765 (20 μM). (E) HEK293T cells were transfected with FLAG-GSDMD-FL, GFP-PLP2 and HA-caspase-1 respectively or together for 24 h, and then the cells were labeled with indicated antibodies. Subcellular localization of FLAG-GSDMD-FL (cyan), GFP-PLP2 (green), HA-caspase-1 (red) and DAPI (blue, nucleus marker) were visualized with confocal microscopy. All results shown are representative of at least three independent experiments.

**S2 Fig. PI staining for** Fig. 1 **I and J.** (A) Vero cells were co-transfected with plasmids encoding GSDMD-FL and caspase-1. At 12 h after transfection, the cells were non-infected or infected with different doses of PEDV for another 24 h, then the cells were performed with propidium iodide (PI) staining analysis. (B) Vero cells were co-transfected with plasmids encoding GSDMD-FL and caspase-1. At 12 h after transfection, the cells were non-infected or infected with PEDV at an MOI of 0.05. At indicated time points after infection, the cells were performed with propidium iodide (PI) staining analysis. Scale bar, 20 μm.

**S3 Fig. The papain-like protease 2 of PEDV inhibits proteasomal degradation of caspase-1.** (A) HEK293T cells were transfected with FLAG-GSDMD-FL and HA-caspase-1 together with increasing amount of MYC-PEDV-PLP2. Cell lysates were analyzed by immunoblotting. (B) HEK293T cells were co-transfected with FLAG-GSDMD-FL together with empty vector or MYC-PEDV-PLP2 then treated with cycloheximide (CHX) (25 μg/ml) for the indicated time points. Cell lysates were analyzed by immunoblotting. (C) Co-immunoprecipitation and immunoblot analysis of extracts of HEK293T cells transfected with FLAG-caspase-1 or FLAG-caspase-1-K134R together with HA-K11 ubiquitin. All results shown are representative of at least three independent experiments.

**S4 Fig. RIG-I was cleaved by caspase-1.** (A) (long exposure of Fig. 4D) HEK293T cells were co-transfected with wild-type (WT) S-MYC-RIG-I or its mutants (D163A, D189A, D194A, D199/202/203A, D209A) together with empty vector or HA-caspase-1. Cell lysates were analyzed by immunoblotting. (B) HEK293T cells were co-transfected with wild-type (WT) h-FLAG-RIG-I or its single, double mutants (D163A, D194A, D234A, D163/194A, D163/234A, D194/234A) together with empty vector or H-HA-caspase-1. Cell lysates were analyzed by immunoblotting. (C) Alignment of the amino acid sequence of RIG-Is from different species. (D) HEK293T cells were co-transfected with wild-type (WT) h-FLAG-RIG-I or its triple mutant (D163/194/234A) together with empty vector or h-HA-caspase-1. Cell lysates were analyzed by immunoblotting. (E and F) IPEC-J2 cells were transfected with RIG-I-WT or RIG-I-D189A, followed by a 12-hour Poly(I:C) stimulation, the indicated gene mRNA levels were quantified by qRT-PCR. All results shown are representative of at least three independent experiments.

**S5 Fig. The papain-like protease 2 of PEDV inhibits proteasomal degradation of caspase-1.** (A) Vero cells were co-transfected with FLAG-GSDMD-FL and HA-Caspase-1, then the cells were infected with PEDV for performed with propidium iodide (PI) staining analysis.

**S6 Fig. PLP2 degrades GSDMD-p30 by removing the K27-linked ubiquitin chains at K275 residue to inhibit pyroptosis.** (A) HEK293T cells were transfected with FLAG-GSDMD-FL together with increasing amount of MYC-PEDV-PLP2. Cell lysates were analyzed by immunoblotting. (B) HEK293T cells were co-transfected with FLAG-GSDMD-p30 and MYC-PEDV-PLP2, then the cells were performed with propidium iodide (PI) staining analysis. All results shown are representative of at least three independent experiments. ****stands for P<0.0001, ***stands for P<0.001, ** stands for P<0.01, * stands for P<0.05 and ns stands for non-significant difference.

**S7 Fig. Papain-like proteases in other CoVs can inhibit pyroptosis.** (A) PI staining for Fig. 7A. HEK293T cells were co-transfected with GSDMD-p30 and PEDV-PLP2 or SARS-CoV-2-PLPro or PDCoV-PLPro, then the cells were performed with propidium iodide (PI) staining analysis. (B) HEK293T cells were co-transfected with FLAG-GSDMD-FL and PEDV-PLP2 or SARS-CoV-2-PLPro or PDCoV-PLPro. Cell lysates were analyzed by immunoblotting.

**S1 Table.** Primers used in this study for the construction of plasmids.

**S2 Table.** qRT-PCR primers used in this study.

**S3 Table.** siRNA sequence for the porcine GSDMD oligonucleotide.

