## Supplemental Table 1 for "Diverse Strategies Utilized by Coronaviruses to Evade Antiviral Responses and Suppress Pyroptosis"

**S1 Table****. Primers used in this study for the construction of plasmids.**

| **Plasmids** | **Forward (5’-3’)** | **Reverse (5’-3’)** |
| --- | --- | --- |
| HA-Caspase-1 | GGAGGCCCGAATTCGGTCGACCATGGCCGATAAGGTGCTGA | CATGTCTGGATCCCCGCGGCCGCTTAATGTCCTGGGAAGAGATAAAAAG |
| FLAG-GSDMD-FL | CAAGCTTGCGGCCGCGAATTCTATGGCATCAGCCTTTGAGAGG | TGCCACCCGGGATCCTCTAGACTAGCAGAGCTGGCTGAGCC |
| MYC-PEDV-PLP2 | ACCTCGGTTCTATCGATTGAATTCGCCACCATGGTATCCACACCTGATGA | GCCAAGCTTCTGCAGGTCGACTTCACAGATCCTCTTCAGAGATGAGTTTCTGCTCGACGACAACATTTGT |
| FLAG-Caspase-1 | AAGGATGACGATGACAAGCTTATGGCCGATAAGGTGCTGAA | CAGGGATGCCACCCGGGATCCTTAATGTCCTGGGAAGAGATAAAAAG |
| FLAG-Caspase-1-K134R | AGGGAGCCTCAGACTTTGCCCCCCAGACATAGC | CAAAGTCTGAGGCTCCCTCTTGGCTCTGAAGAC |
| HA-Caspase-1-C285A | TTATCCAGGCTGCACGTGGTGAGAAGCAAGGGG | ACGTGCAGCCTGGATAATGATCACCTTGGGTT |
| MYC-PEDV-PLP2-C113A | GATAATAACGCATGGGTTAATGTTACATGTTTACAATTACA | ACCCATGCGTTATTATCTGTGGTTTTAAGCACACG |
| MYC-PEDV-PLP2-H272A | TGTCGGCGCATATACTGTTTTTGATCATGACACTGGTAT | CAGTATATGCGCCGACAACACCGTTACCATTA |
| MYC-PEDV-PLP2-D285A | TGGTGCATGCAGGAGATGCTTTTGTACCGGGT | ATCTCCTGCATGCACCATACCAGTGTCATGATC |
| HA-Caspase-1-C285A-K134R | AGGGAGCCTCAGACTTTGCCCCCCAGACATAGC | CAAAGTCTGAGGCTCCCTCTTGGCTCTGAAGAC |
| S-MYC-RIG-I | GGAGGCCCGAATTCGGTCGACTATGACAGCAGAGCAGCGGC | CATGTCTGGATCCCCGCGGCCGCCTACTCAAGGTTGCCCATTCCC |
| S-MYC-RIG-I-D163A | CTCAGATCAGCTAAGGAGAATTGGCCTAAAACTTTG | CTCCTTAGCTGATCTGAGGAGGCATTCAACCAT |
| S-MYC-RIG-I-D189A | GATGGTAGCTAAAGGTGCAGAAGATGTTAAAATGAA | GCACCTTTAGCTACCATCCACAGTTCACTGAACC |
| S-MYC-RIG-I-D194A | CAAAGGTGCAGAAGCTGTTAAAATGAAAGATCTTGAGGATGACG | ACAGCTTCTGCACCTTTGTCTACCATCCACAGT |
| S-MYC-RIG-I-D199/202/203A | AAAGCTCTTGAGGCTGCTGAAATGAAAACTTGTGATGTACAGATTT | AGCAGCCTCAAGAGCTTTCATTTTAACATCTTCTGCACCT |
| S-MYC-RIG-I-D209A | AACTTGTGCTGTACAGATTTTCTACAAGGAAGAACC | TCTGTACAGCACAAGTTTTCATTTCGTCATCCT |
| S-MYC-RIG-I-1-189aa | GGAGGCCCGAATTCGGTCGACTATGACAGCAGAGCAGCGGC | CATGTCTGGATCCCCGCGGCCGCCTAGTCTACCATCCACAGTTCACTGAA |
| S-MYC-RIG-I-190-943aa | GGAGGCCCGAATTCGGTCGACTAAAGGTGCAGAAGATGTTAAAATGA | CATGTCTGGATCCCCGCGGCCGCCTACTCAAGGTTGCCCATTCCC |
| h-FLAG-RIG-I-D163A | CAGCAAAGGAAAACTGGCCCAAAACTTTGAAAC | GCCAGTTTTCCTTTGCTGATCTGAGAAGGCATTCCACC |
| h-FLAG-RIG-I-D194A | AAGCAGTTGAAACAGAAGATCTTGAGGATAAGA | CTTCTGTTTCAACTGCTTTTATACCTTTCTCTACAATCCACAGT |
| h-FLAG-RIG-I-D234A | CAGAAGTGTCTGCTACAAACTTGTACAGCCCATTTAAACC | TGTAGCAGACACTTCTGAAGGTGGACATGAAT |
| h-Caspase-1 | CAAGCTTGCGGCCGCGAATTCCATGGCCGACAAGGTCCTG | CCTCTAGAGTCGACTGGTACCTTAATGTCCTGGGAAGAGGTAGAAA |
| FLAG-GSDMD-p30 | AAGGATGACGATGACAAGCTTATGGCATCAGCCTTTGAGAGG | CAGGGATGCCACCCGGGATCCTCAGTCTGACTGGAACTTCAGGTG |
| FLAG-GSDMD-p30-K103R | AGGACAGGGGAGATTCTCCGGCGGGGCCGCGGT | GAGAATCTCCCCTGTCCTGGGGCTGCCAGCTCC |
| FLAG-GSDMD-p30-K177R | GACCCACAGACAGGAGGGCTCGGGCCAGTTTGC | CCCTCCTGTCTGTGGGTCCGCGTGACCTCCACC |
| FLAG-GSDMD-p30-K275R | AGCACCTGAGATTCCAGTCAGACGGGCCCGCGGAG | ACTGGAATCTCAGGTGCTCAGAGGGGAAGCTCA |
| FLAG-GSDMD-FL-K275R | AGCACCTGAGATTCCAGTCAGACTGAGGATCCC | ACTGGAATCTCAGGTGCTCAGAGGGGAAGCTCA |
