## Supplemental Table 2 for "Diverse Strategies Utilized by Coronaviruses to Evade Antiviral Responses and Suppress Pyroptosis"

**S2 Table. qRT-PCR primers used in this study.**

| **Name** | **Forward (5’-3’)** | **Reverse (5’-3’)** |
| --- | --- | --- |
| Porcine Caspase-1 | CAGGAGTCCTCGAACTCTCCACAG | GGCTCTGAAGACGCAGGCTTAAC |
| Porcine IFN-α | ACCTTCCAGCTCTTCAGCACAGA | TCCTGCATGACACAGGCTTCCA |
| Porcine IFN-β | CCTGGAACAGTTGCCTGGGACT | TCTGCTGGAGCATCTCGTGGAT |
| Porcine ISG15 | ATGGGTAGGGAACTGAAGGT | CAGACGCTGCTGGAAGG |
| Porcine OAS1 | GCCTGTGATTCTGGACCCGGCTGA | CGACACCTTCCAGGATCCCACCG |
| PEDV-S | CGGTTTGTTGGATGCTGTC | AATAAAGAATACGCTGAATGGC |
| β-actin (IPEC-J2) | TGCGGCATCCACGAAACTAC | AGGGCCGTGATCTCCTTCTG |
| GAPDH (Vero) | CACTGAGGACCAGGTTGTGTCCTGTGAC | TCCACCACCCTGTTGCTGTAGCCAAATTC |
| human IFN-β | AAACTCATGAGCAGTCTGCA | AGGAGATCTTCAGTTTCGGAGG |
| human ISG15 | AGATCACCCAGAAGATCG | TGTTATTCCTCACCAGGATG |
| GAPDH (THP-1) | GCCTTCCGTGTCCCCACTG | CGCCTGCTTCACCACCTTC |
