## Supplemental Table 3 for "Diverse Strategies Utilized by Coronaviruses to Evade Antiviral Responses and Suppress Pyroptosis"

**S3 Table. siRNA sequence for the porcine GSDMD oligonucleotide**

| **Name** | **Forward (5’-3’)** | **Reverse (5’-3’)** |
| --- | --- | --- |
| Si-GSDMD#1 | CCCUUCUACUUCCAUGACATT | UGUCAUGGAAGUAGAAGGGTT |
| Si-GSDMD#2 | GCUGAAUGAAACCCAGCAUTT | AUGCUGGGUUUCAUUCAGCTT |
| Si-GSDMD#3 | CUGCACGUGUGCACUUUAUTT | AUAAAGUGCACACGUGCAGTT |
